# EC-Bench: A Benchmark for Enzyme Commission Number Prediction

**DOI:** 10.1101/2025.06.25.661207

**Authors:** Saeedeh Davoudi, Christopher S. Henry, Christopher S. Miller, Farnoush Banaei-Kashani

## Abstract

Enzymes are proteins that catalyze specific biochemical reactions in cells. Enzyme Commission (EC) numbers are used to annotate enzymes in a four-level hierarchy that classifies enzymes based on the specific chemical reactions they catalyze. Accurate EC number prediction is essential for understanding enzyme functions. Despite the availability of numerous methods for predicting EC numbers from protein sequences, there is no unified framework for evaluating and studying such methods systematically. This gap limits the ability of the community to identify the most effective approaches for enzyme annotation. We introduce EC-Bench, a benchmark for EC number prediction, consisting of 1) an initial representative set of existing methods (including homology-based, deep learning, contrastive learning, and language model methods), 2) existing and novel accuracy and efficiency performance metrics, and 3) selected datasets to allow for comprehensive comparative study. EC-Bench is open-source and provides a framework for researchers to not only compare among existing methods objectively under uniform conditions, but also to introduce and effectively evaluate performance of new methods in a comparative framework. To demonstrate the utility of EC-Bench, we perform extensive experimentation to compare the existing EC number prediction methods and establish their advantages and disadvantages in a variety of prediction tasks, namely “exact EC number prediction”, “EC number completion” and (partial or additional) “EC number recommendation”. We find wide variation in the performance of different methods, but also subtle but potentially useful differences in the performance of different methods across tasks and for different parts of the EC hierarchy.

## 1 Introduction

Proteins are biopolymers primarily made up of 20 canonical amino acids. A primary role of proteins is to function as enzymes, which serve as catalysts to accelerate nearly all chemical reactions occurring within cells. [1] The Enzyme Commission (EC), a four-level hierarchical classification system (e.g., 1.1.1.1), not only provides EC numbers as broadly used annotations for enzymes in genome analyses but also links these enzymes to the specific chemical reactions they catalyze within metabolic pathways. [2] Definitive enzyme annotation relies on experimental methods, which are time-consuming and struggle to keep pace with the rapidly increasing volume of newly discovered protein sequences. To address these challenges, researchers have explored various computational approaches and predictive models to infer the EC numbers for enzymes.

Traditional homology-based approaches (i.e., methods that rely on sequence similarity for EC prediction) [3, 4, 5, 6] predict enzyme functions by aligning the enzyme sequence to those in annotated databases, presuming that similar sequences are likely to have similar evolutionary history and thus similar functions. Machine Learning (ML) models, a technology that empowers computers to learn from data and make predictions without being explicitly programmed for specific tasks, offer new ways to predict EC numbers for enzymes by recognizing more complex patterns in the sequences rather than relying solely on sequence alignment [7, 8, 9, 10, 11]. Moreover, recognizing that protein sequences share structural similarities with text, some researchers have turned to language models [12]. These models, originally designed for Natural Language Processing (NLP), have been adapted for EC number prediction by interpreting protein sequences in a manner analogous to text [13, 14, 15, 16]. Finally, techniques like contrastive learning have been leveraged to improve EC prediction accuracy. By training models to distinguish between similar and dissimilar examples, contrastive learning models [17, 18] have pushed the boundaries of predictive performance in this domain.

Effective benchmarks have a long history of promoting and measuring advancements in protein data analysis. However, currently there is no comprehensive benchmark for EC prediction methods. CASP (The Critical Assessment of Protein Structure Prediction) [19] is a benchmark that aims to evaluate the accuracy of protein structure prediction methods by providing an open-source platform for the research community to perform comparative empirical studies on structure prediction methods. Inspired by CASP, CAFA (Critical Assessment of Function Annotation) [20] was designed to enable assessment of computational methods dedicated to predicting Gene Ontology (GO) terms. While historically helpful for assessing protein annotation prediction, CAFA is primarily a benchmark *dataset* and competition rather than a comprehensive “one-stop shop” benchmark platform (like our proposed benchmark), which in addition to a benchmark dataset offers implementation of a representative set of function annotation models in the field, and a collection of complementary metrics for fair evaluation. Such a benchmark platform allows researchers to develop and integrate new models into the benchmark system and seamlessly perform an objective comparative assessment versus state-of-the-art function annotation models. Moreover, such a benchmark platform enables studying all existing function annotation models in an objective way to settle their advantages and disadvantages for the benefit of the research community and the practitioners alike. TAPE (Tasks Assessing Protein Embeddings) [21] and FLIP (Fitness Landscape Inference for Proteins) [22] introduce benchmarks specifically designed to evaluate the effectiveness of transformer-based protein language models across multiple biologically relevant tasks. ProteinGym [23] provides a collection of benchmarks specifically crafted for protein fitness prediction and design. However, no AI-ready benchmark like ProteinGym or FLIP exists for the task of predicting EC numbers based on protein sequence. CARE [24] is a recent benchmark for EC number prediction, covering both protein-to-EC number and reaction-to-EC number mapping. Protein sequences and their associated EC numbers used in CARE were sourced from SwissProt, the expertly curated subset of UniProtKB [25]. While CARE evaluates the accuracy of a few models, it does not incorporate a broad range of models available in the field, limiting its comprehensiveness in benchmarking and learning from comparisons across diverse approaches.

Despite the availability of a wide array of computational methods for predicting EC numbers from protein sequences, there is no benchmarking tool that allows for systematic evaluation of these methods under uniform conditions. The current EC prediction landscape is fragmented, with different models being evaluated on different datasets, employing varied performance metrics, and utilizing different data preprocessing techniques.

This lack of standardization makes it difficult to directly compare the efficacy of different approaches, as differences in dataset selection, evaluation metrics, and preprocessing steps can all significantly impact the reported results. Consequently, lack of a benchmark for fair evaluation of the EC prediction methods makes it challenging for researchers to determine which models are truly effective for enzyme annotation.

Here, we introduce EC-Bench, an open-source benchmarking tool that includes a representative set of 10 existing EC number prediction methods selected based on a comprehensive literature review (Table 1), while providing researchers with a platform to incorporate and evaluate new methods against existing techniques under uniform assumptions. EC-Bench includes a carefully curated standardized dataset and performance metrics that enable thorough evaluation of accuracy and efficiency for EC number prediction methods. We conduct extensive experimentation using EC-Bench to compare existing EC number prediction methods, highlighting their advantages and disadvantages across varied tasks

**Table 1:**
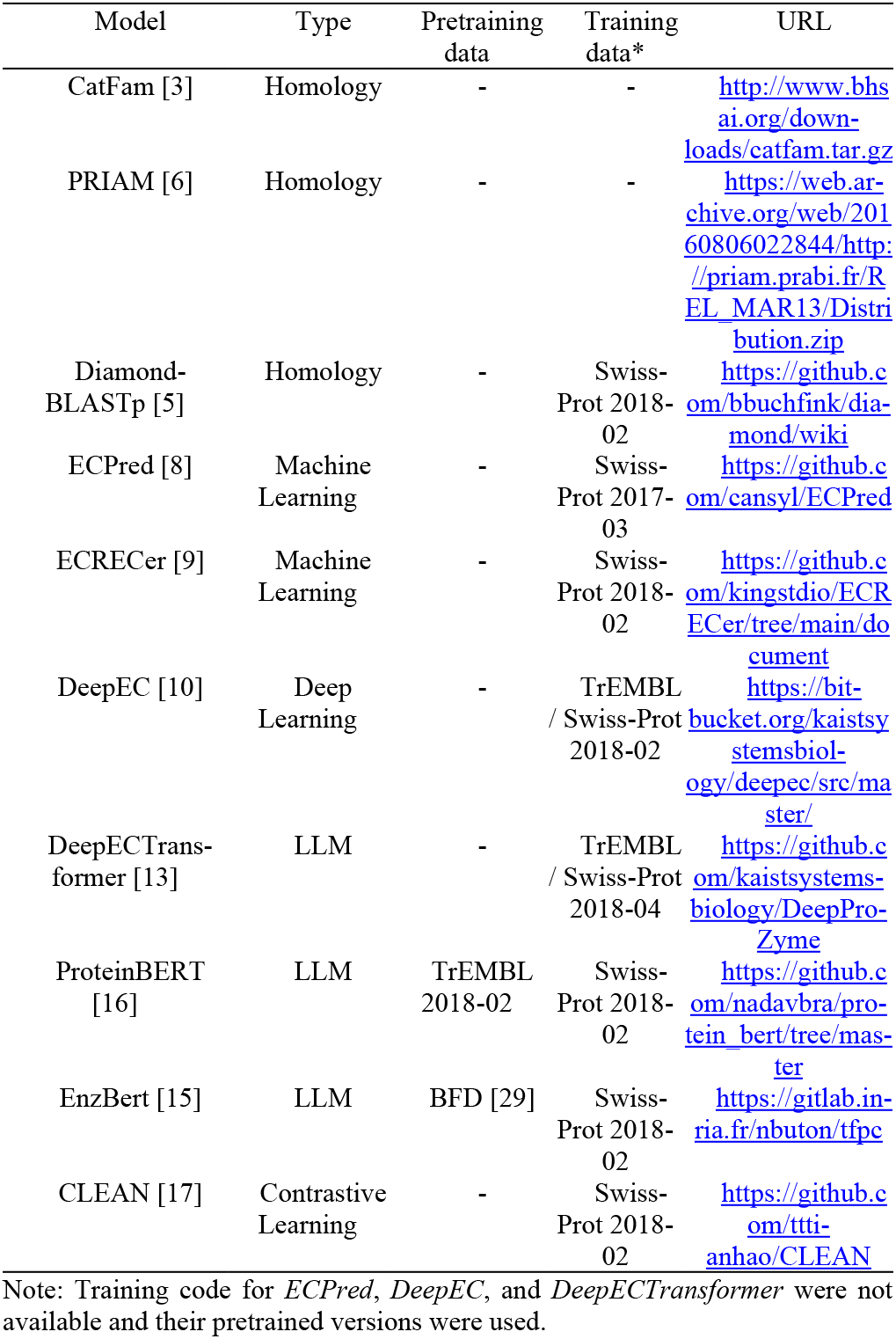
Benchmark Models in EC-bench.

## 2 Materials and Methods

To address the gap in evaluating EC number prediction methods, we developed EC-Bench (Figures 1, 2), a benchmark framework designed for consistent and thorough assessment of EC number prediction. EC-Bench consists of three key components: *Benchmark Dataset (Figure 1), Model Pretraining and Training*, and *Evaluation Tasks (Figure 2)*, each designed to ensure objective and consistent evaluation.

**Figure 1.**
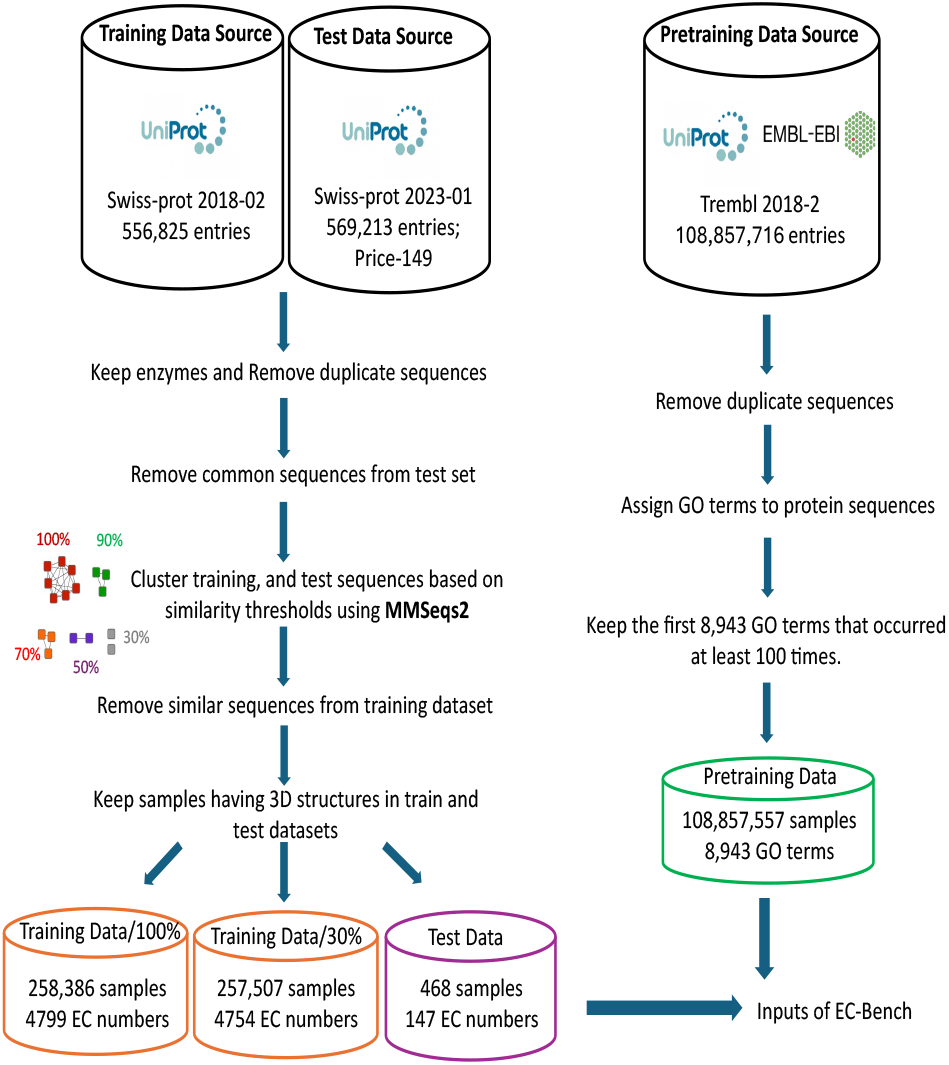
Overview of benchmark datasets and data preprocessing pipeline. The left side illustrates the preprocessing steps for generating training and test datasets, while the right side shows the preparation of the pretraining dataset. The final output includes two training sets (for 30% and 100% similarity thresholds across test and training data), one test set, and one pretraining set, all structured for use within the EC-Bench framework.

**Figure 2.**
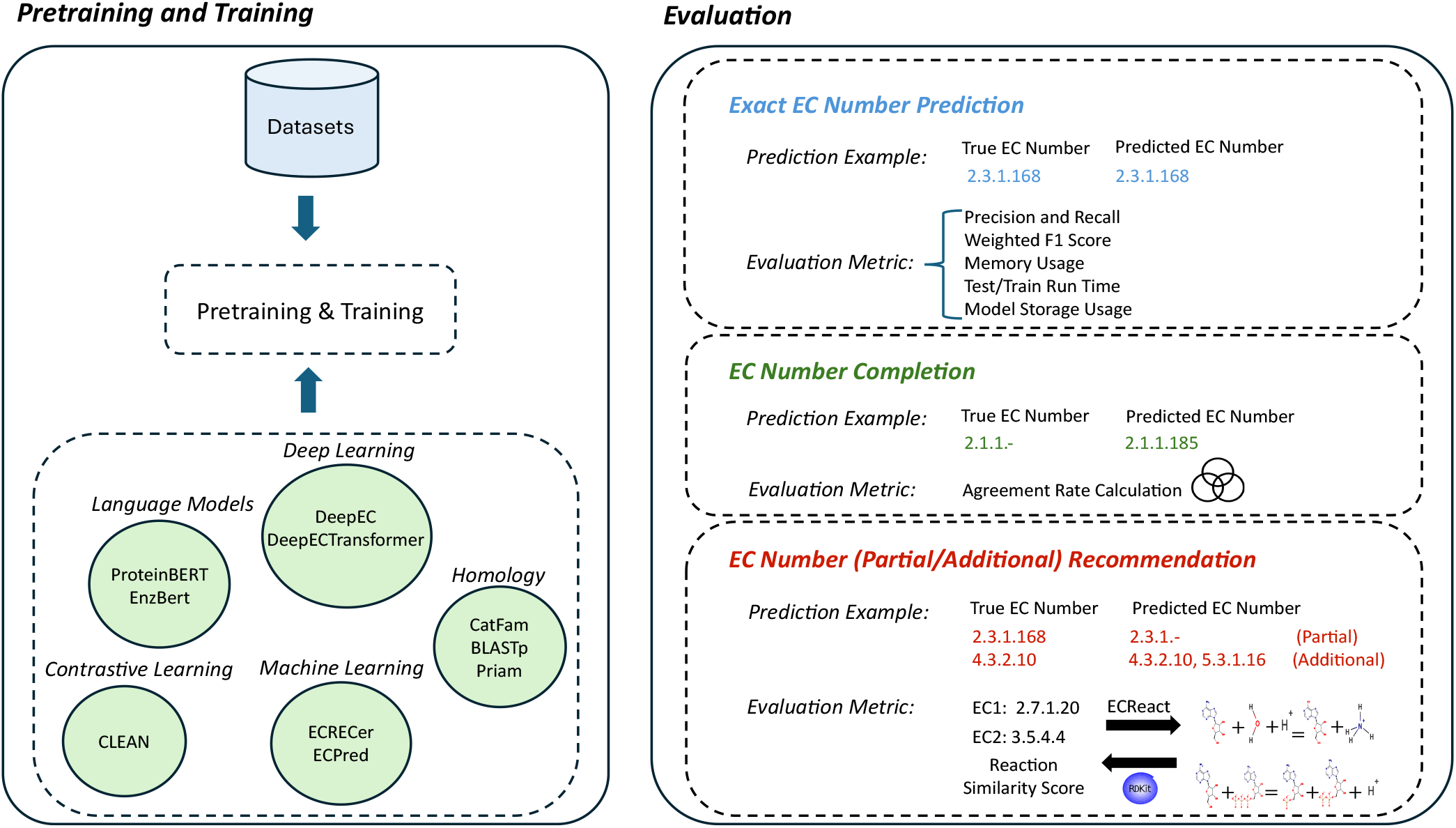
Pretraining, training, and evaluation tasks of EC-Bench. First, models are pretrained and trained using datasets curated by EC-Bench (left side). Then, the trained models are evaluated across three tasks (right side). For each evaluation task, an example prediction and the corresponding evaluation metrics are illustrated.

### 2.1 Benchmark Dataset

#### 2.1.1 Dataset construction

We constructed pretraining, training, and test datasets suitable for evaluation of all models (Figure 1) Protein sequences and their corresponding EC numbers were primarily sourced from UniProtKB, a high-quality database. UniProtKB consists of two major components: Swiss-Prot, a manually curated dataset with high annotation reliability, and TrEMBL, which contains automatically annotated sequences. Given the importance of using well-curated and standardized datasets for benchmarking, we carefully selected different subsets of these datasets for pretraining, training, and testing:

- *Pretraining Data*: Extracted from *TrEMBL version 2018-02*, this dataset provides a large-scale, automatically annotated collection of protein sequences. It is particularly useful for models such as ProteinBERT [16] that require extensive pretraining on diverse sequence representations and functions before fine tuning.
- *Training Data*: Derived from *Swiss-Prot version 2018-02*, this manually curated dataset ensures high-quality EC number annotations. This version was chosen to include the models DeepEC [10] and ECPred [8], which do not provide capabilities for retraining, but were originally trained on this same version.
- *Test Data*: 1) *Swiss-Prot version 2023-01* was used as the primary test dataset, as it represents a recent version available at the time of evaluation. This dataset ensures that models are tested on the latest curated sequences, reflecting real-world conditions; 2) *Price-149*, a challenging dataset of 149 experimentally curated bacterial protein sequences. Price-149 was previously used in the ProteInfer [11] and *CLEAN* [17] studies and is known to contain cases where homology-based methods struggle, making it useful for evaluating model robustness.

#### 2.1.2 Data Preprocessing

The training and test datasets were preprocessed to improve data quality and enhance model generalization. Highly similar sequences could lead to inflated performance metrics where models memorize patterns instead of learning more complex meaningful sequence-function relationships. On the other hand, highly similar sequences with different functions could also be useful to models that can learn from these subtle differences. Therefore, we filtered out similar sequences at different similarity thresholds to evaluate the effect of retaining similar sequences across models. Additionally, since our goal is to benchmark enzyme function prediction, we excluded non-enzyme sequences and kept only those with available 3D structural models to support future structure-aware modeling. The preprocessing steps are as follows:

1. *Removing Non-Enzyme Proteins:* Non-enzyme sequences (those without annotated EC numbers) were removed from training and test datasets.
2. *Removing Duplicate Sequences:* Removing duplicates from within pretraining, training, and test datasets prevents bias in training and evaluation.
3. *Similar Sequence Filtering across test and training data:* This step ensures that models are evaluated on their ability to generalize beyond sequences they have already encountered. Sequences were filtered out of the training dataset at similarity thresholds of 100%, 90%, 70%, 50%, and 30%. This filtering ensured that identical (100%) or similar (other thresholds) sequences to the test data were not present in the training data, preventing data leakage and overfitting. We used MMseqs2 [26] to cluster all training and test data protein sequences at each threshold. We then check all test sample clusters and remove any training data sequences found in the same cluster.
4. *Keeping Enzymes with 3D Structures:* With the recent advent of accurate 3-D protein structure prediction methods, we anticipate structure-informed EC prediction will become a growing trend. Only enzymes with available AlphaFold [27] structural models were kept in training and test datasets, facilitating EC-Bench’s future applicability to structure-aware function prediction.
5. *Integration of Gene Ontology (GO) Terms:* Finally, we retrieved GO terms associated to protein sequences in the pretraining dataset from the EMBL-EBI database [28]. GO terms provide a broader functional context for models to learn from, capturing molecular functions, biological processes, and cellular components.

Table 2 shows the final dataset sizes across different similarity thresholds.

**Table 2:** Size of datasets per each similarity threshold.

| Similarity threshold | Pretraining data size | Training data size | Test data size |
| --- | --- | --- | --- |
| 30% | 108,857,557 | 257,507 | 468 |
| 50% | 108,857,557 | 257,773 | 468 |
| 70% | 108,857,557 | 257,986 | 468 |
| 90% | 108,857,557 | 258,321 | 468 |
| 100% | 108,857,557 | 258,386 | 468 |

Note: Similar sequences are only removed from the training data.

### 2.2 EC Prediction Model Selection

EC-Bench includes 10 diverse models selected based on their availability, compatibility with the dataset, and prominence in the field. These models span a range of all major approaches introduced in the field, including homology-based approaches, deep learning, contrastive learning, and language models (Table 1).

Additionally, we incorporated two ensemble learning methods in EC-Bench. By incorporating ensemble learning, EC-Bench evaluates whether combining methodologically diverse models can enhance EC number prediction accuracy. Two ensemble approaches are:

1. *Majority Voting:* A straightforward approach where the final prediction is determined by the most frequently predicted EC number.
2. *Stacking:* A more advanced ensemble method in which the predictions from a subset of models on a validation dataset are used as input features for a meta-model, a Multi-Layer Perceptron classifier (see parameters in Table S1). This meta-model is then trained on input features to predict EC numbers on the test dataset.

EC-Bench also provides users with the flexibility to integrate and evaluate their own models within the benchmarking framework, and to compare them objectively to existing state-of-the-art approaches.

### 2.3 Evaluation Tasks

The EC-Bench workflow (Figure 2) begins with training (and pretraining for some models) and proceeds to evaluate models on query proteins. EC-Bench employs three tasks to evaluate models (shown in different colors in Figure 2). These tasks are designed to evaluate models in terms of accuracy, EC number completion, and the ability to recommend partial EC numbers or additional (i.e. multi-function) EC numbers.

#### 2.3.1 Exact EC Number Prediction

This task measures how accurately models predict exact EC numbers. Since EC numbers follow a structured classification system, model performance is evaluated at increasing levels of specificity (levels 1 through 4).

##### 2.3.3.1 Accuracy Measurement

For accuracy measurement, we consider precision, recall, and weighted F1 score to compare different models in EC-Bench. Precision measures the proportion of correctly predicted positive labels out of all labels predicted as positive by the model. Recall, also known as sensitivity or true positive rate, measures the proportion of correctly predicted positive labels out of all actual positive labels. The weighted F1 score is a metric that combines both precision and recall into a single number, providing a balanced measure of a model’s accuracy. Unlike the regular F1 score, the weighted F1 accounts for the imbalance in the dataset by considering the support (i.e., the number of true instances) of each class (i.e., distinct EC numbers). The formula for the F1 score for a single EC number class is:

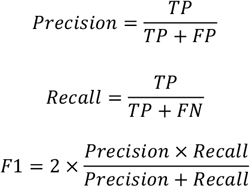

Where TP is True Positives,

FP is False Positives, and FN is False Negatives. The weighted F1 score is then calculated by:

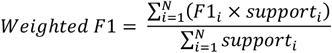

where F1_i_ is the F1 score for EC number *i*, support_i_ is the number of true instances for EC number *i*, and *N* is the total number of distinct EC numbers. A high precision indicates that when the model predicts a positive EC number, it is correct most of the time, high recall means that the model predicts correct EC Numbers for most of the true positives, which is crucial in scenarios where missing a positive case is costly, and a high weighted F1 score indicates that a model performs well across both precision and recall while accounting for EC number class imbalances in the dataset.

##### 2.3.1.2 Optimizing EC Number Prediction Thresholds

The EC number prediction task is formulated as a multi-label classification problem, as a single enzyme can be associated with multiple EC numbers corresponding to different catalytic activities (i.e. multi-function). In multi-label classification tasks, deciding how to convert predicted probabilities into discrete class assignments is crucial for achieving accurate and meaningful predictions. A standard binary classification model typically applies a fixed threshold (e.g., 0.5) to determine whether a class should be assigned to a given instance. However, this approach may not be optimal for EC number prediction, where some classes may require more conservative thresholds while others benefit from more inclusive thresholds. For example, consider a neural model that uses a sigmoid activation function at the output layer and optimizes binary cross-entropy loss across all labels. The sigmoid activation function maps the output of each neuron to a probability value between 0 and 1, making it appropriate for multi-label classification tasks where each label is predicted independently. For a given input *x*, the sigmoid function is defined as:

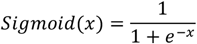

This output represents the predicted probability that a specific EC number is present for the given input. Binary cross-entropy loss is commonly used in multi-label classification to evaluate the difference between the predicted probabilities and the actual binary labels (1 for presence, 0 for absence). Together, the sigmoid activation and binary cross-entropy loss allow a model to predict the presence of multiple EC numbers for a single enzyme.

However, if the model assigns probabilities of 0.45 to EC number A, 0.48 to EC number B, and 0.52 to EC number C, a uniform threshold of 0.5 would only assign EC number C, potentially missing relevant classifications. Conversely, if a protein has multiple functional annotations with probabilities just below 0.5, a strict thresholding approach may fail to capture all relevant functions. To mitigate these challenges, we introduce two complementary strategies as a wrapper solution:

1. *Learning Class-Specific Thresholds (“Learnt Models”):* Instead of applying a fixed threshold across all classes, EC-Bench learns an optimal threshold for each EC number (class). This is done by maximizing the F1 score for each class empirically, evaluating threshold values from 0 to 1 in increments of 0.01. During prediction, these learned thresholds are used to classify an instance into a particular EC number. If none of the predicted probabilities exceed their respective learned thresholds, the model selects at least the class with the highest probability. This approach is particularly useful for rare EC numbers, where setting a higher threshold may prevent false positives, while more common EC numbers may benefit from a lower threshold to avoid missing relevant classifications.
2. *Adaptive Decision-Making Based on Probability Distributions (“Regular Models”):* An alternative strategy is to use a more flexible, adaptive thresholding method, where the model makes decisions based on the relative magnitudes of predicted probabilities rather than fixed values.

- If the highest predicted probability for a given class exceeds a highconfidence threshold (0.8), the model assigns that class.
- Additional classes are considered only if their probabilities exceed a secondary threshold (0.5), allowing for multi-label classification where appropriate.

These balances ensure that the model selects confident predictions, while allowing multiple assignments when necessary.

##### 2.3.1.3 Performance Efficiency

We also measure the computational efficiency of different models during both training and inference. We record the peak amount of runtime memory usage during model training, the total disk storage space required for the trained model file (model size), and the total duration required to train the model and to generate predictions on the test dataset during inference (run times). Collectively, these metrics help evaluate the suitability and efficiency of the various models for deployment and/or scalability in a variety of environments and use cases.

#### 2.3.2 EC Number Completion

EC number completion is an important task in enzyme function prediction because many enzymes in biological databases are only partially annotated. These are represented with dashes in place of the unknown levels (e.g., *1*.*1*.*-*. *-*), indicating that while the enzyme’s broad function is known (or inferred), finer details are still missing. Completing EC numbers allows researchers to infer more precise enzyme functions, which is essential for understanding metabolic pathways. In this evaluation task, we assess the ability of models to fill in missing levels of incomplete EC numbers (“-”) in the test data at any level.

As we do not have the true completed EC numbers to directly validate the predictions, to evaluate models on this task we introduce a metric named *agreement rate*. The agreement rate measures the proportion of incomplete EC numbers for which most of the models converged on the same completed prediction. Specifically, it is calculated as the percentage of cases where more than half of the models returned the same EC number as their top prediction for a given incomplete entry. This metric serves as an indicator of inter-model consistency and highlights how often models independently arrive at a shared decision when faced with uncertain or under-specified input. A high agreement rate suggests that the model is making consistent and potentially confident predictions. Conversely, a low agreement rate may indicate high variability in model behavior, reflecting either uncertainty in the input data or diverse model strategies for EC number completion.

#### 2.3.3 Partial/Additional EC Number Recommendation

In many real-world biological scenarios, there are enzymes whose exact functions are unknown. Even partial EC number recommendations may be valuable for inferring participation in metabolic pathways or to suggest experimentally testable hypotheses. In contrast to exact EC Number Prediction (section 2.3.1), this task evaluates the ability of models to generalize to new EC numbers (*partial*, as full EC number may be absent from the training data) and to suggest *additional* EC numbers for multi-function enzymes.

Because the true recommended EC numbers are not explicitly available for validation, and experimental validation is not feasible at scale, we need a task-specific novel evaluation metric to assess how biologically relevant the model’s recommendations are. As EC numbers represent chemical reactions, we chose to evaluate predictions by comparing the similarity between the reactions of the existing true and predicted EC numbers using RDKit [30]. Reaction similarity scores between two EC Numbers range from 0 to 1.

Enzymatic reactions and their corresponding EC numbers were previously extracted from major databases and merged into a unified dataset called ECReact [31]. However, we found the list of EC numbers in ECReact is incomplete. In cases where an EC number is incomplete or missing, we ignore the fourth level of the EC number and randomly sample EC numbers that share the first three levels. We compute the reaction similarity as the average similarity between the sampled EC numbers and the true EC number. If no EC numbers exist that share the first three levels, we progressively reduce the specificity by ignoring additional levels until we can calculate an average reaction similarity. For cases where there are multiple predicted EC numbers or multiple true EC numbers, we take the maximum similarity score among the EC numbers.

To compare models together across all samples, we report weighted similarity score for each model:

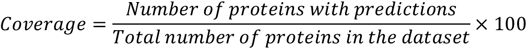

Higher coverage means the model can make predictions for a larger fraction of proteins.

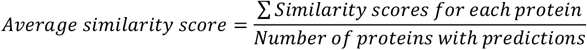

Average similarity score measures the functional similarity between predicted and true EC numbers.

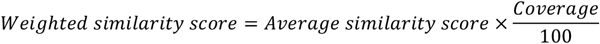

Weighted similarity score combines accuracy (similarity score) with coverage, ensuring models that predict for more proteins are not penalized unfairly.

## 3 Results and Discussion

In the following subsections, we present a detailed analysis of each evaluation task described above, integrating these findings and discussing their implications for enzyme annotation and functional prediction. We evaluate models under two sequence similarity thresholds: *100% similarity* (where test sequences may have closely related training counterparts) and *30% similarity* (where test sequences are more dissimilar from the training data). This comparison highlights the trade-offs between homology-based, deep learning-based, contrastive learning, and ensemble methods, offering insights into their strengths and weaknesses in different sequence similarity scenarios and for different parts of the EC hierarchy.

### 3.1 Results of Exact EC Number Prediction

#### 3.1.1 Accuracy Measurement

To evaluate exact EC number prediction, we analyze weighted F1 scores, precision, and recall across the four EC hierarchy levels.

##### 3.1.1.1 Weighted F1 Score

Overall, there was a surprisingly high level of variability in model performance. On the test dataset, weighted F1 scores at EC level 1 ranged from 0.164 to 0.906 at the 100% similarity threshold, and from 0.164 to 0.895 at the 30% threshold. At EC level 4, scores ranged from 0.074 to 0.639 (100%) and from 0.074 to 0.524 (30%), highlighting the increasing difficulty of finegrained classification as sequence similarity in the training data decreases. *EnzBert-learnt* achieved the highest weighted F1 scores at levels 1, 2, and 3 for both similarity thresholds on the test dataset (Figure 3a, Figure 3b). At 100% similarity, its F1 scores for level 1, 2, and 3 are 0.873, 0.777, and 0.692, while at 30% similarity, they remain stable at 0.895, 0.794, and 0.702, respectively. At level 4 on the 100% similarity test set, the homology-based *BLASTp* slightly outperforms deep learning models, achieving a weighted F1 score of 0.502 compared to *EnzBert-learnt*’s F1 score of 0.475 (Figure 3a). However, at 30% similarity, *EnzBert-learnt* surpasses BLASTp, achieving an F1 score of 0.463 compared to *BLASTp*’s score of (0.431) (Figure 3b). Across both similarity thresholds, other models performed nearly as well as the top-performing models.

**Figure 3.**
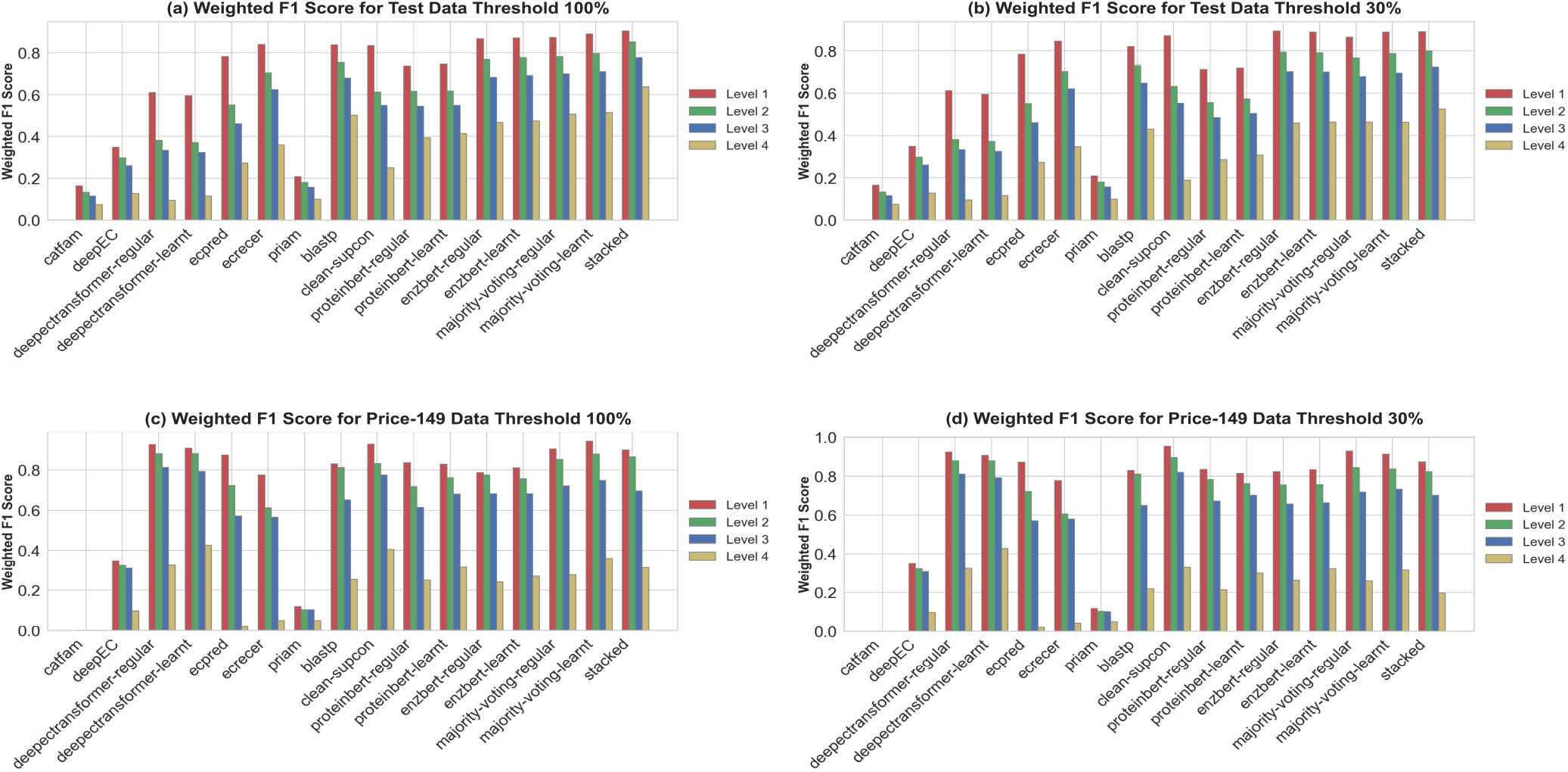
Weighted F1 scores for exact EC number prediction across all models and test sets(a) Weighted F1 scores for the test set at 100% data similarity threshold. (b) Weighted F1 scores for the test set at 30% data similarity threshold. (c) Weighted F1 scores for the Price-149 test set at 100% data similarity threshold., (d) Weighted F1 scores for the Price-149 test set at 30% data similarity threshold.

On the more challenging Price-149 dataset, *DeepECTransformer-learnt* achieves the highest F1 score at level 4 (0.427), followed by *CLEAN* (0.406) (Figure 3c). However, because we could not separately train *DeepECTransformer*, this model may have had data leakage. Except for *CLEAN* and *DeepECTransformer*, all models exhibit a drop in their weighted F1 scores when moving from the test dataset to the out-of-distribution Price-149 dataset.

Collectively, these results suggest that while commonly used homology-based approaches like *BLASTp* perform well when sequence similarity to proteins in the training set is high, language models such as *EnzBert-learnt* generalize better across diverse enzyme families and are more robust when test sequences are distantly related to training sequences. Many models, like the contrastive-learning-based *CLEAN* generalize better to out-of-distribution data than homology-based models.

##### 3.1.1.2 Precision and Recall Trade-offs

Interesting model-type-specific patterns emerged when comparing precision and recall across the different test datasets. *BLASTp* consistently achieves the highest precision across all levels at both test set similarity thresholds (Figure S1a, S1b). However, at level 4, its precision drops when sequence similarity to the training data decreases, from 0.612 (100% similarity) to 0.527 (30% similarity) (Figure S1a vs. S1b). *CLEAN* also declines in precision at level 4, dropping from 0.488 (100%) to 0.308 (30 %) (Figure S1a vs. S1b). The LLM-based model *EnzBert-learnt* demonstrates better precision stability across the test datasets, with only a minor drop at level 4 from 0.511 (100%) to 0.49 (30%) (Figure S1a vs. S1b). It also maintains the highest recall across all levels, with minimal change from 100% to 30% similarity (Figure S2a, S2b). This suggests that this LLM-based model captures functional relationships beyond strict sequence similarity. On Price-149, *CLEAN* maintains its superiority at level 4 for both precision (Figure S2c, S2d) and recall (Figure S3c, S3d) at 100% and 30% similarity thresholds. Overall, the trade-off between precision and recall is different across different model types: while homology-based approaches offer high precision but suffer from recall loss when similarity decreases, deep learning models maintain stable recall.

##### 3.1.1.3 Impact of Ensemble Learning

When sufficient training data is available, the stacked model outperforms all other models. With the 100% similarity threshold, the weighted F1 is 0.906 at level 1 and 0.639 at level 4, (Figure 3a). At 30% similarity, the stacked model’s performance declines slightly, with an F1 score of 0.892 at level 1 and 0.524 at level 4 (Figure 3b). However, on the Price-149 dataset, the stacked model fails to outperform other models (Figure 3d), likely due to its reliance on training distributions that differ from Price-149. This highlights the need for ensemble models to integrate more diverse data during training to generalize effectively to challenging datasets.

The simpler majority voting model does not always outperform other models, as its performance depends on the collective predictions of individual models. When most models predict an incorrect EC number, the voting mechanism reinforces this error, leading to suboptimal performance. In such cases, individual models that make correct predictions can achieve higher accuracy than the majority voting approach.

##### 3.1.1.4 Model Comparison Using EC Hierarchical Plots

To provide a more detailed comparison between models, EC-Bench generates interactive HTML sunburst plots (See Supplementary Material) that visualize the F1 scores for each EC number at each hierarchical level. For each pair of models, two sunburst plots allow side by side comparison, along with a third plot highlighting the differences in their F1 scores across all EC numbers. These visualizations offer an intuitive understanding of where in the EC hierarchy models agree and where they differ in prediction quality.

For example, when comparing *BLASTp* and *CLEAN* at 100% similarity threshold, we observed that BLASTp outperforms CLEAN on EC numbers associated with fewer training samples *(e*.*g*., 4.2.3.127). However, BLASTp tends to perform worse when predicting more frequent and diverse EC numbers such as 2.7.7.7. In contrast, when comparing *BLASTp* and *EnzBert*, the behavior of the two models is more similar overall, perhaps reflecting the complementary nature of sequence similarity-based and language-model-based approaches. Additionally, a comparison between *CLEAN* and *EnzBert-learnt* shows that *EnzBert-learnt* consistently performs better on EC numbers that have either a large number of training samples (e.g., 2.5.1.-) or very few samples (e.g., 4.2.3.127). This highlights the ability of large language models to generalize both in data-rich and data-scarce scenarios, unlike *CLEAN*, which tends to struggle on these same examples where data is highly imbalanced.

#### 3.1.2 Performance Efficiency

We compared the computational efficiency of all methods in training and inference. Inference run times varied over several orders of magnitude. *ECRECer* and *ProteinBERT* are among the fastest methods at inference (2 and 17 seconds on test data; Table 4). ECRECer and BLASTp require minimal memory, making them especially beneficial for large, high-similarity datasets. However, *BLASTp*’s dependence on close homology causes a considerable decline in performance at the 30% threshold (Figures 3.b, 3.d), and ECRECer relies on Evolutionary Scale Modeling (ESM) embeddings [32] which means it needs extra preprocessing before making predictions. In contrast, *EnzBert* and *ProteinBERT* demand more computational resources (Table 3) but may justify this overhead by remaining more resilient across varying similarity levels (Figures 3.a–3.d). Notably, *ProteinBERT* stands out for its relatively fast inference among deep learning/LLM models, thanks to a smaller parameter count (Table 4). The contrastive-learning model *CLEAN* offers memory efficiency yet requires extensive training (5160 epochs at 100% similarity and 1875 at 30%). *CLEAN* also requires ESM embeddings, which can add additional time to the overall process. This time for computing embeddings was not included in the run times reported in Tables 3 and 4. Despite this, *CLEAN*’s inference phase is relatively quick once the model is fully trained.

**Table 3:** Training run time, storage and memory usage for models.

| Model | Memory usage (GiB) | Model size (MiB) | Run time per epoch* | Total training run time |
| --- | --- | --- | --- | --- |
| ECRECer | 6.1 | 5000 | - | 1327s |
| CLEAN | 8.2 | 4 | 30s | 2d (5160 epochs (100%), 1875 epochs (30%)) |
| ProteinBERT | 40 | 394 | 150s | 3.5d (81 epochs) |
| EnzBert | 17 | 1640 | 1d | 15d (15 epochs) |
| Diamond-BLASTp | 1.9 | 681 | - | 17s |

Note: Includes models trained from scratch. s: seconds, d: days, \*on a RTX6000-40GB GPU.

**Table 4:**
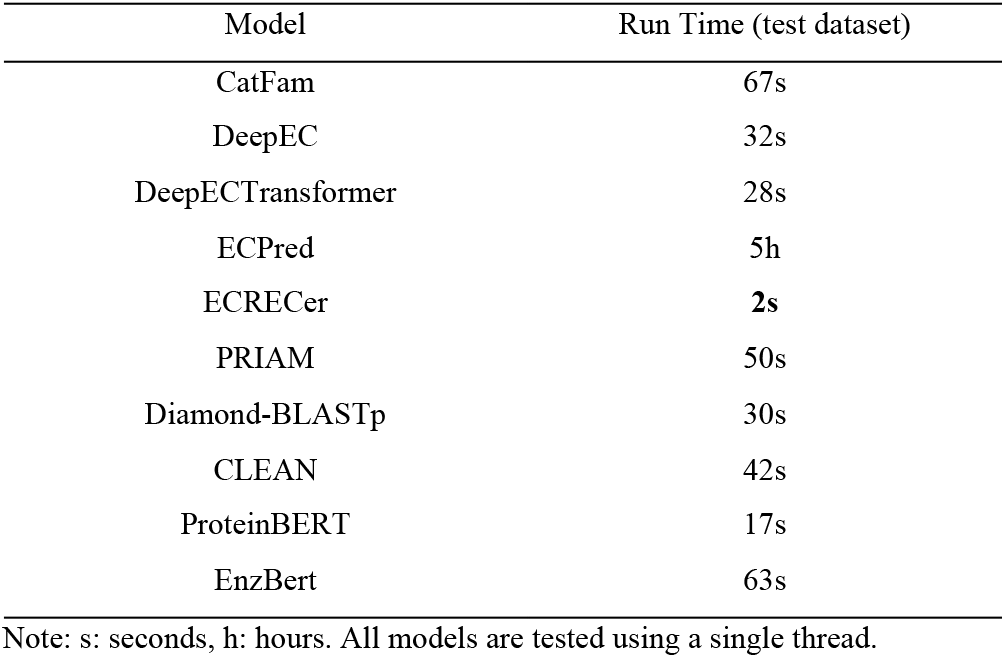
Test run time for all models.

### 3.2 Results of EC Number Completion

Many enzymes with existing annotations do not have complete 4-level EC numbers. In this section, we evaluate the ability of models to complete EC numbers for proteins in the test data that do not have full 4-level EC Number annotations, asking them to fill in missing parts of enzyme classifications. As by definition we do not know the correct 4-level classification for this task, for evaluation purposes we compare both the number of completions made, and the agreement rate among models. *CLEAN* completes notably more EC numbers than other models at level 4 and in total (Figure S4). Agreement rate highlights a model’s alignment with the collective prediction trend, offering insight into which models are more consistent with consensus, and which tend to diverge in their predictions (Figure 4). Notably, *CLEAN* had the highest number of participations in majority agreements, suggesting that it is both accurate when evaluated on proteins with full annotations (Section 3.1.1), and consistent with the broader ensemble when evaluated on incomplete annotations, making it a strong candidate for EC number completion in ambiguous or partially annotated cases.

**Figure 4.**
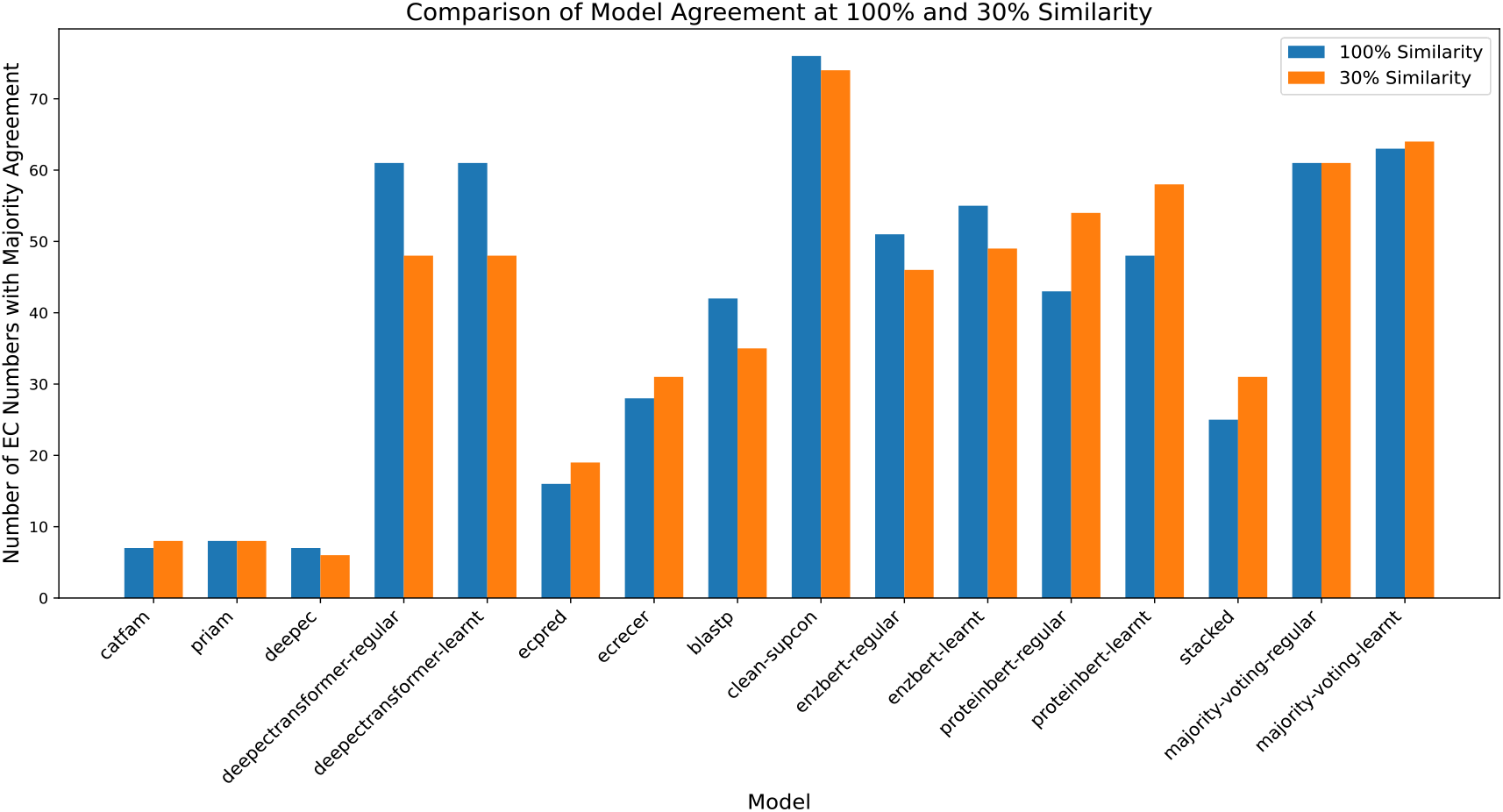
Model participation in majority EC number agreements. For each model, the number of EC numbers in which it participated in a majority agreement is reported. An EC number is considered to have majority agreement if more than half of the models predicted it for the same protein. This analysis reflects how consistently each model aligns with the consensus across the dataset, highlighting differences in model conformity and prediction behavior.

### 3.3 Results of Partial EC Number Recommendation

We next evaluated the ability of the models to recommend new partial EC numbers (levels 1-3) for EC numbers that exist in the test data but were never seen in the training data (Figure S5). Compared to other models, the *EnzBert-learnt* model predicts more closely related EC numbers across all three levels at the 30% similarity threshold. This suggests that *EnzBertlearnt* excels in handling more diverse and less similar sequences, likely due to its ability to generalize well in more complex data scenarios. However, at the 100% similarity threshold, the top three models across all levels are *majority voting-regular, CLEAN*, and *ProteinBERT-learnt*. The shift in model performance between thresholds highlights the varying strengths of different models depending on the nature of the input data.

Partial EC number predictions might still have value if they are “close” to correct (i.e., enzymes are predicted to perform a chemical reaction similar to the true reaction). Therefore, we also evaluated how similar predicted partial EC numbers were to the true reaction for this task. At the 100% similarity threshold, almost all models perform nearly equally well (Figure 5a), predicting reactions similar to the truth for many of the unseen EC numbers. However, at the 30% similarity threshold, there is more variability, with some models demonstrating better generalization capabilities when challenged with lower sequence similarity (Figure 5b).

**Figure 5.**
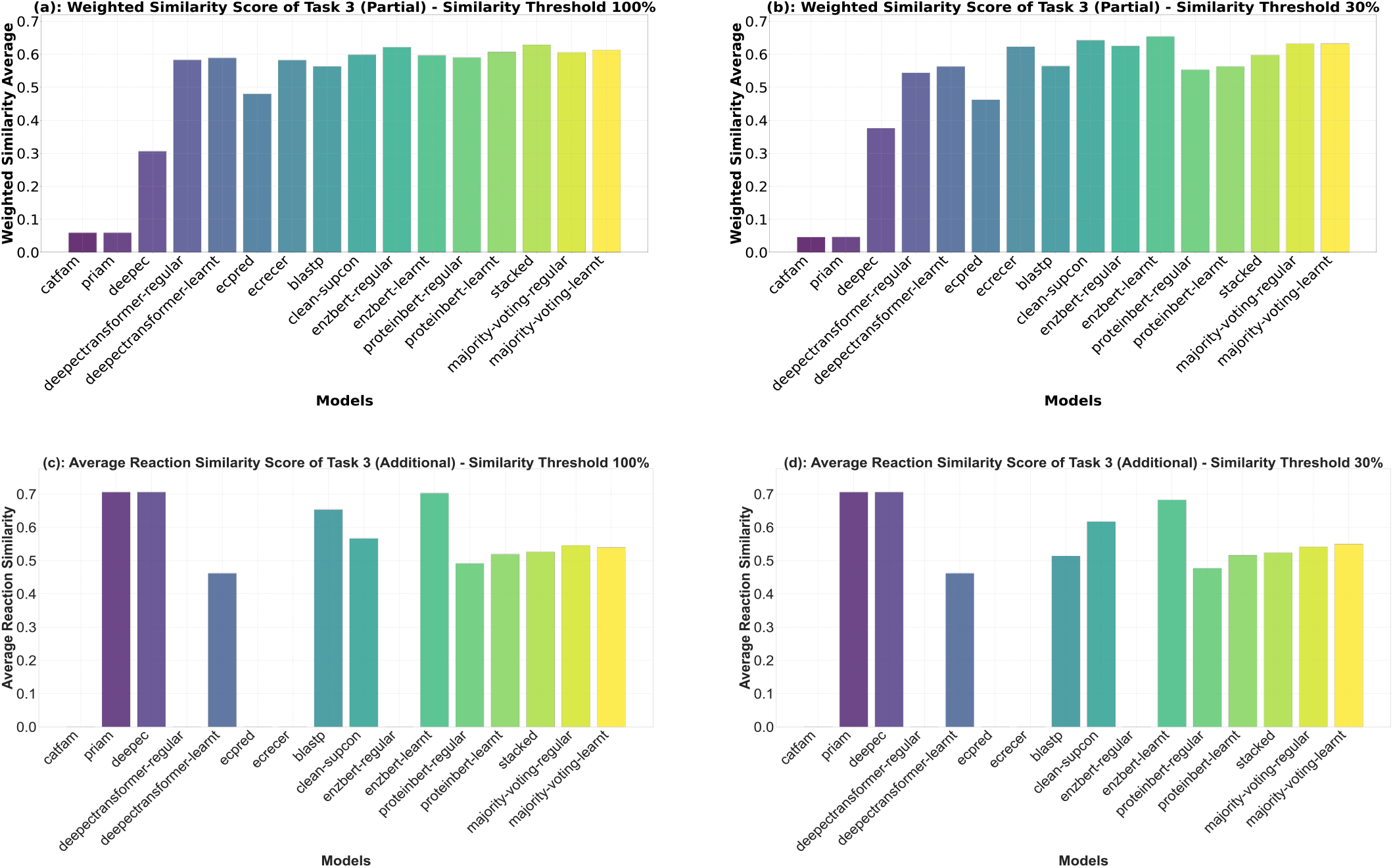
Reaction similarity scores for models on EC number recommendation tasks. Panels (a) and (b) illustrate the weighted reaction similarity score for partial EC number recommendation at 100% and 30% sequence similarity thresholds, respectively, highlighting the ability to predict partial EC numbers at varying levels of specificity for EC numbers in the test data that are unseen in the training data. Panels (c) and (d) present the average reaction similarity scores for additional EC number recommendation at 100% and 30% sequence similarity thresholds, respectively, highlighting the ability to predict additional (multi-function) EC numbers with good reaction similarity to existing annotations.

### 3.1 Results of Additional EC Number Recommendation

Some enzymes can have multiple functions, and models should be able to recommend additional functions when appropriate. We first counted cases where models correctly predicted the annotated function, but also provided additional EC Number predictions (Figures S6a and S6b). *Majority voting* models have the tendency to predict additional EC numbers due to ties in voting; however, excluding these, *EnzBert-learnt* (at the 30% threshold) and *ProteinBERT-learnt* (at 100%) predict the most additional EC numbers. If correct, language models thus might be useful for suggesting multiple or additional enzymatic functions. It is also possible that these new EC numbers could represent corrections to the annotated function.

Under the assumption that additional new functions should be metabolically related to anntoted ones (Figure S7), we computed a reaction similarity score between existing and additional new annotations (Figures 5c and 5d). Many models suggest additional functions with average reaction similarity scores similar to the distribution of known multi-function enzymes. Whether our assumption holds and these are indeed accurate recommendations of new additional functions will need experimental validation.

## 4 Conclusion

In this work, we introduced EC-Bench, a benchmark for evaluating EC number prediction methods. EC-Bench provides an end-to-end solution, covering data collection, preprocessing, model training, and evaluation, while enabling objective comparisons of diverse predictive models. Through extensive experiments on 10 representative models, including homology-based, deep learning, contrastive learning, and language models, we demonstrated the platform’s ability to compare models under standardized conditions. By providing a unified framework, EC-Bench facilitates the development and evaluation of new methods, supporting advancements in computational enzyme annotation.

Homology-based methods have been the annotation method of choice for decades. We found *BLASTp*’s precision is high when dealing with well-characterized enzyme families at high similarity, but a limitation of any benchmark is that almost all existing database annotations were assigned via homology. Despite this, language models and threshold-optimized methods excel in more diverse or novel datasets. With these caveats, minimizing false positives favors homology-based approaches, but when discovering new enzymes or functional variants, the higher recall of language models is advantageous.

For some models, their agreement rates, average reaction similarity scores, and weighted similarity scores are higher under the 30% similarity threshold compared to the 100% threshold. This is somewhat counterintuitive, as more similar sequences available in training should in theory equate to better performance in testing. However, some models may also generalize better with a training set with slightly less redundancy. This behavior warrants further investigation in future work.

## Acknowledgements

This work used the computing resources at the Center for Computational Mathematics, University of Colorado Denver, including the Alderaan cluster, supported by the National Science Foundation award OAC-2019089.

## Data availability

The data underlying this article and full benchmark implementation with documentation are available in GitHub, at https://github.com/dsaeedeh/EC-Bench.

