## Supplementary material for "EC-Bench: A Benchmark for Enzyme Commission Number Prediction": sup-material.pdf

### Supplementary Tables

| Parameter | Value | Description |
|---|---|---|
| Hidden layer size | 100 | One hidden layer containing 100 neurons. |
| epochs | 200 | Number of iterations (epochs) for training the model. |
| alpha | 0.001 | Regularization parameter (L2 penalty) to prevent overfitting. |
| Learning rate | Adaptive, initial value: 0.001 | Adjusts the learning rate dynamically, reducing it when loss plateaus. |

Table S1. Parameters of the MLP classifier used for stacking training.

| Parameter | Value / Description |
|---|---|
| Base model | ProtBert-BFD pretrained on protein sequences from BFD |
| Dropout on (CLS) | 0.2 |
| Batch size | 2 |
| Accumulation step | 16 |
| Learning rate | $1.0 \times 10^{-5}$ |
| Optimizer | Adam ($\beta_1 = 0.9$, $\beta_2 = 0.999$) |
| Lr scheduler | Lr (epoch) = 0.8 × Lr (epoch − 1) |
| Number epochs | 15 |

Table S2. EnzBert finetuning parameters.

| Parameter | Value / Description |
|---|---|
| Pretraining dataset | ~106 million protein sequences from UniRef-90 |
| Model variants | Supports sequence lengths of 128, 512, and 1024 tokens |
| Architecture style | Inspired by BERT, with 6 transformer blocks, combining local (convolutional/FC) and global (attention) pathways |
| Global attention | Uses linear-complexity global-attention layers enabling support for very long sequences |
| Parameter count | ~16 million parameters, significantly smaller than similar models |
| Pretraining tasks | Masked language modeling + Gene Ontology (GO) annotation prediction |
| Activation function | GELU |
| Optimizer | Adam |
| Number of epochs | ~6.4 passes over the dataset (28 days of training) |
| Hardware/training speed | ~280 protein records per second on an Nvidia Quadro RTX 5000 GPU |
| Fine-tuning strategy | Freeze all layers + train classification head (40 epochs), then unfreeze full model (40 additional epochs), finishing with 1 epoch at longer sequence length |
| Fine-tuning duration per task | ~14 minutes on a single GPU, across 9 benchmarks |

Table S3. ProteinBERT training parameters.

### Supplementary Figures

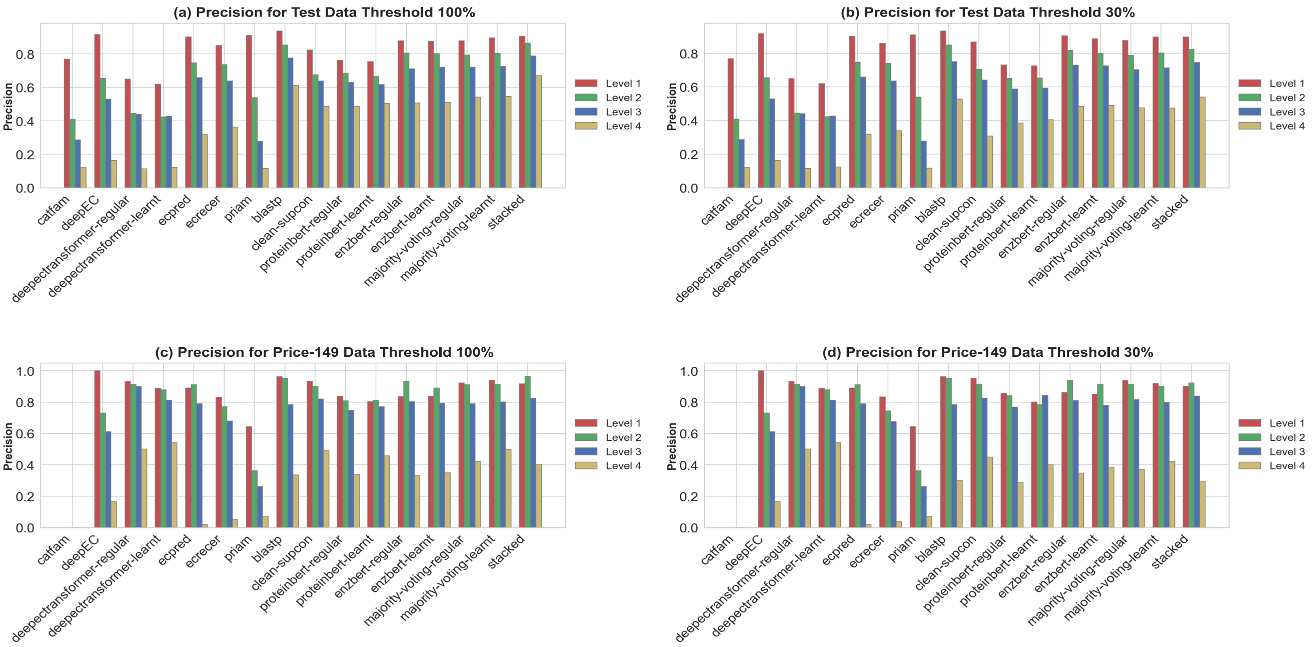


Figure S1: Precision for exact EC number prediction across all models and test sets(a) precision for the test set at 100% data similarity threshold. (b) precision for the test set at 30% data similarity threshold. (c) precision for the Price-149 test set at 100% data similarity threshold. (d) precision for the Price-149 test set at 30% data similarity threshold.

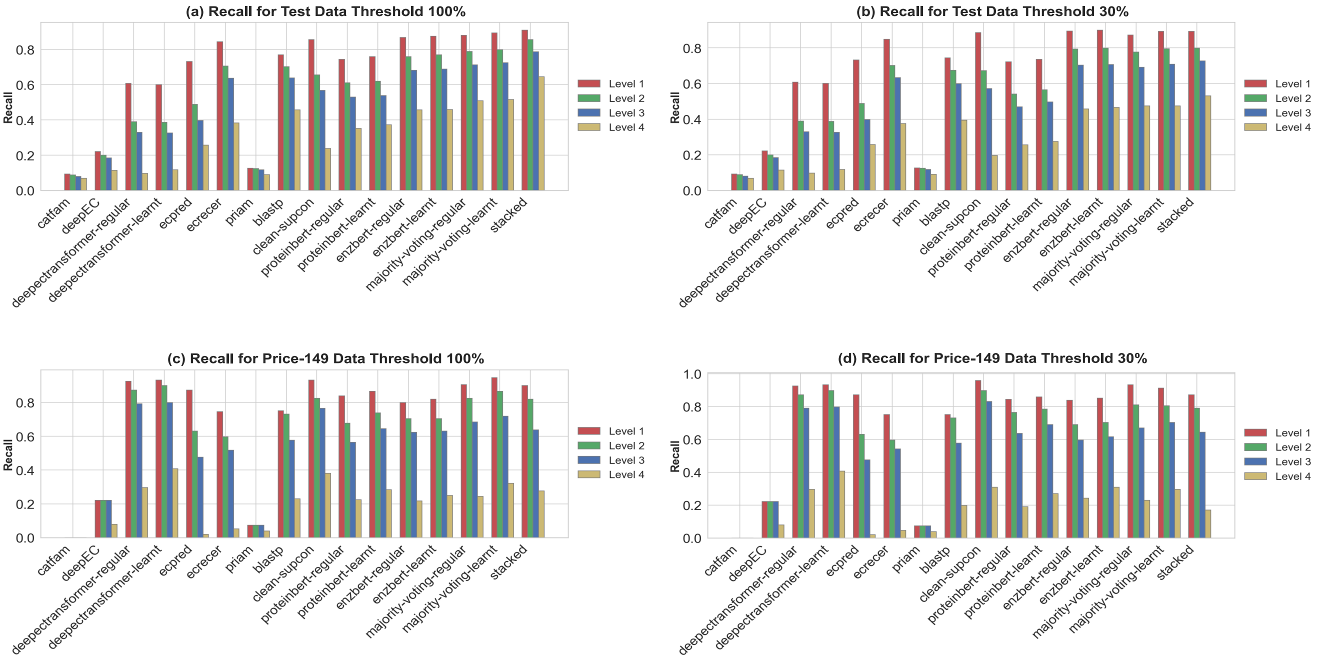


Figure S2: Recall for exact EC number prediction across all models and test sets(a) recall for the test set at 100% data similarity threshold. (b) recall for the test set at 30% data similarity threshold. (c) recall for the Price-149 test set at 100% data similarity threshold. (d) recall for the Price-149 test set at 30% data similarity threshold.

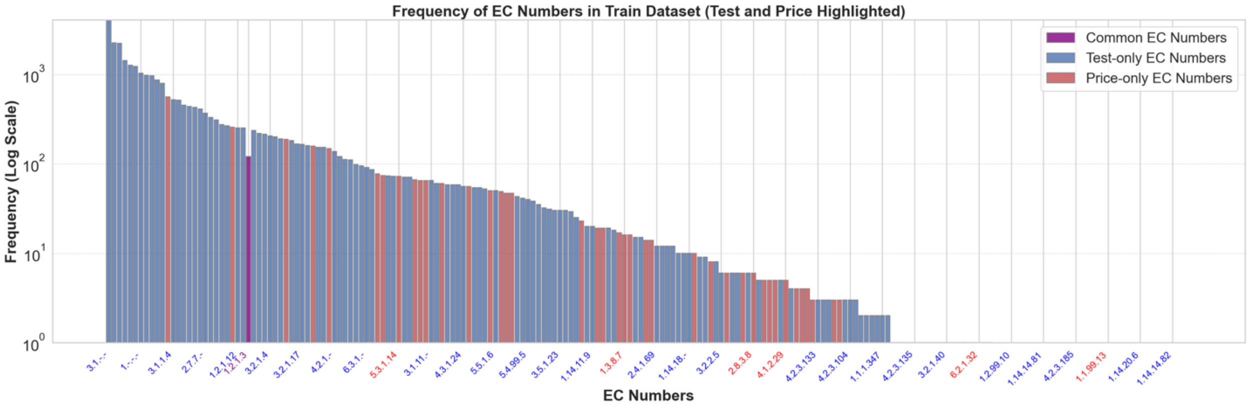


Figure S1: Frequency of EC Numbers in the Train Dataset with Test and Price-149 Datasets Highlighted: The plot displays the frequency of EC numbers from the train dataset, divided into three categories. Purple bars represent EC numbers that are common between the test and Price-149 datasets. Blue bars indicate EC numbers that are exclusive to the test dataset, while red bars represent EC numbers exclusive to the Price-149 dataset. The y-axis is displayed on a logarithmic scale to emphasize differences in frequency, and key EC numbers (common and every 5th label) are highlighted on the x-axis for clarity.

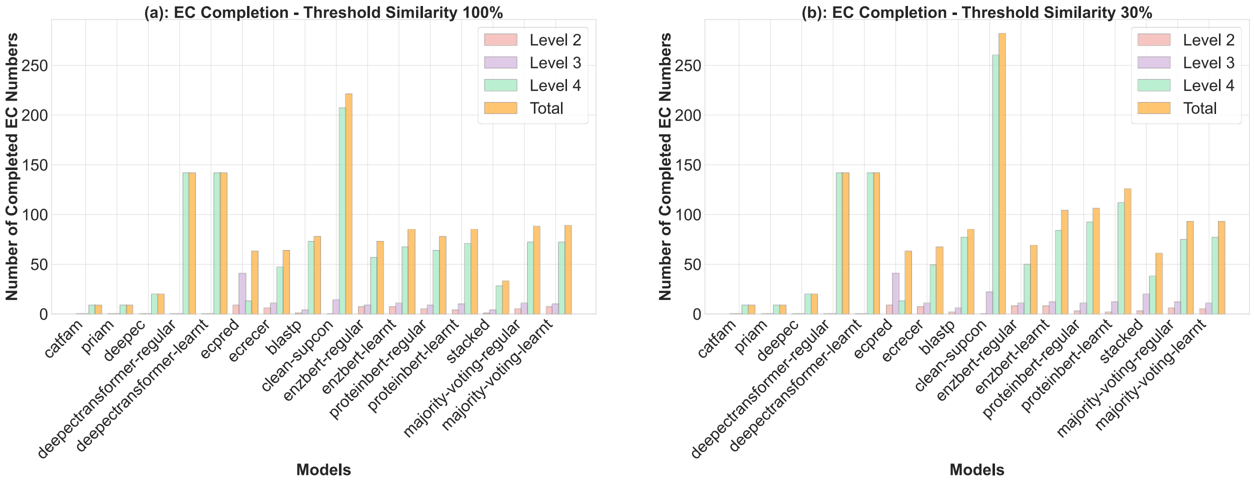


Figure S2: Comparison of models in completing EC numbers under different sequence similarity thresholds. (a) shows the number of EC numbers completed at 100% similarity, while (b) presents the results at 30% similarity. Bar heights represent completions at levels 2, 3, and 4, with an additional bar indicating the total completions per model. This visualization highlights each model’s ability to provide more specific EC annotations as well as its overall performance across multiple levels of detail.

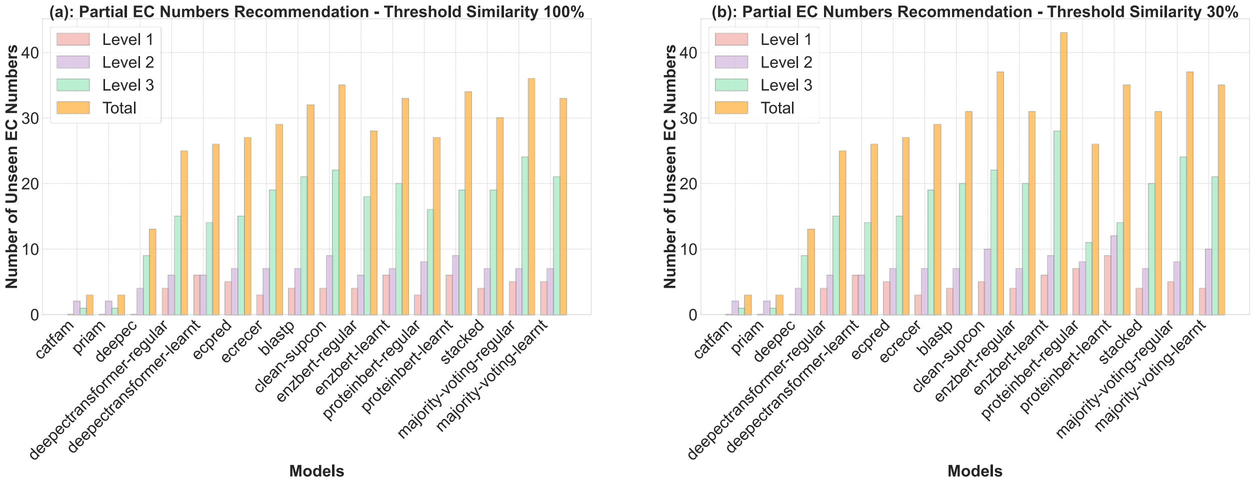


Figure S5: Partial EC number recommendation performance across models at different sequence similarity thresholds. (a) shows the number of unseen EC numbers correctly recommended by each model at levels 1, 2, and 3 for sequences with a 100% similarity threshold, while (b) presents the results for sequences with a 30% similarity threshold. Bars are grouped to represent the number of EC numbers correctly recommended at each level, with an additional bar for the total recommendations. This visualization highlights the ability of models to recommend unseen EC numbers partially.

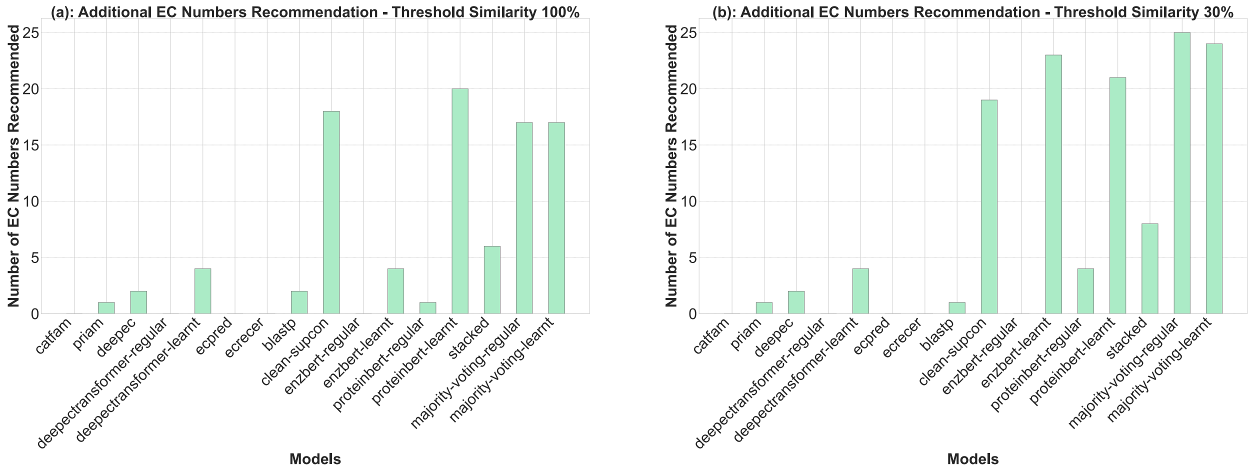


Figure S6: Comparison of “additional” EC number recommendation across models at two sequence similarity thresholds. (a) displays the number of EC numbers recommended by each model for sequences at a 100% similarity threshold, whereas (b) shows the results for sequences at a 30% similarity threshold. The analysis focuses on the models' ability to provide supplementary EC number predictions beyond the true annotations, offering insights into their capacity to identify related enzymatic functions. Each bar represents the total additional EC numbers predicted per model.

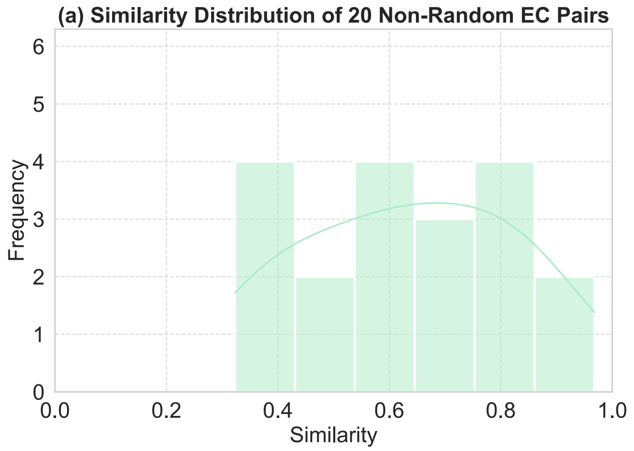


Figure S7: Distribution of pairwise reaction similarity scores for (a) 20 pairs of EC numbers from the same multi-functional enzymes, and (b) 20 pairs of randomly selected EC numbers. The results show that EC numbers from multi-functional enzymes tend to have higher similarity scores (closer to 1), while random EC pairs typically show low similarity (closer to 0).

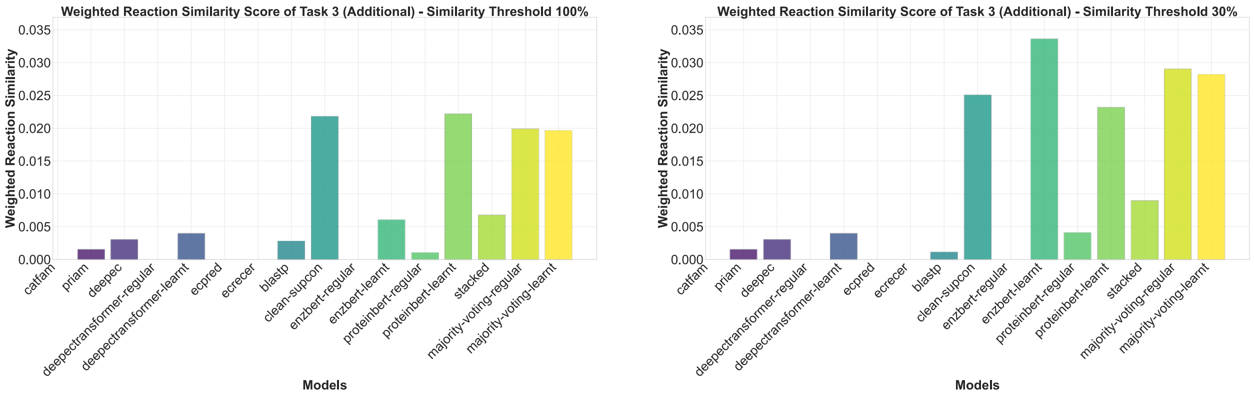


Figure S8. Weighted similarity score of models on “additional” EC number recommendation tasks at 100% and 30% sequence similarity thresholds, respectively. Each bar represents the performance of a model under the respective task and threshold.

### Supplementary Methods

#### Overview of EC number prediction models

In this section, we offer overviews of models used in this benchmark with publicly available software implementations. These models were selected based on their code availability, compatibility with the data, and their prominence in the literature.

1. ***PRIAM***
   In 2003, PRIAM [1] was introduced and later updated to version 2 in 2018. It functions as a profile-based model that predicts enzyme functions by employing a classification scheme found in the ENZYME database [2]. The model conducts a homology search between each protein sequence and its respective profile, resulting in a list of matches. Following this, a predefined enzyme-specific rule is applied to each ENZYME entry to determine if it fulfills the criteria, ultimately generating a list of predicted enzymes. PRIAM undergoes testing on the Swiss-Prot database [3], showcasing high precision and recall in its performance.

2. ***CatFam***
   CatFam [4], proposed in 2008, utilizes protein sequences to construct a profile database of function annotations, achieving high precision in this regard. It also excels in generating hypotheses, although with a slightly lower precision but better recall. Throughout the process of creating the database, researchers carefully monitor the adjustable false-positive rate to maintain an acceptable precision level for each profile.

3. ***BLASTp***
   BLASTp (Basic Local Alignment Search Tool for proteins) [5] is a widely used bioinformatics tool designed to compare an amino acid query sequence against a protein sequence database, identifying regions of local similarity between sequences. This tool plays a crucial role in predicting the function of unknown proteins by aligning them with known sequences, thus inferring potential functional and evolutionary relationships. BLASTp's ability to quickly search and align large databases makes it invaluable for tasks such as identifying homologous proteins, annotating genes, and understanding protein structure and function. The tool is accessible through the NCBI (National Center for Biotechnology Information) website [6], which provides both an online interface and downloadable software for more extensive local analyses.

4. ***ECPred***

In 2018, the ECPred [7] model was proposed, offering distinct models for predicting each level of the EC number, from level 0 to level 4. This model amalgamates the outcomes of three separate predictors; each trained on the Swiss-Prot database at their respective levels. Although ECPred outperforms previous models, it is ineffective for handling multi-functional enzymes, and its performance in predicting 4-level EC numbers is suboptimal.

5. ***DeepEC***

Further progress in 2018 gave rise to the DeepEC [8], which can predict 4-level EC numbers and multi-functional enzymes. DeepEC consists of three independent but coherent Convolutional Neural Networks (CNNs): CNN-1 for enzyme and non-enzyme predictions, CNN-2 for predicting EC numbers up to level 3, and CNN-3 for predicting EC numbers up to level 4. Additionally, a homology-based predictor, BLASTp, is integrated into the model as a fallback in case CNN-2 and CNN-3 fail to predict EC numbers accurately.

6. ***ECRECer***

In 2022, a new model called ECRECer [9] has emerged in this domain. ECRECer is based on multiagent dual-core learning and boasts an exciting capability to predict the fourth level of incomplete EC numbers. ECRECer can do three tasks: 1) enzyme vs non-enzyme classification task, 2) single or multi-functional enzyme classification task, and 3) EC number prediction task. Moreover, it can provide recommendations for the top 20 EC numbers with the highest scores for an unknown enzyme.

7. ***CLEAN***

CLEAN (Contrastive Learning-Based Enzyme Annotation) [10] is an innovative model designed to enhance the accuracy of enzyme annotation by leveraging contrastive learning techniques. Unlike traditional models that rely primarily on sequence alignment or feature extraction alone, CLEAN distinguishes between similar and dissimilar protein sequences through contrastive learning. CLEAN's algorithm operates by first encoding protein sequences into high-dimensional representations using a neural network architecture. It then applies contrastive learning, a technique that involves creating pairs of similar and dissimilar sequences based on known functional annotations. The model is trained to minimize the distance between representations of sequences that share the same EC number while maximizing the distance between those that do not. This training process enables CLEAN to learn fine-grained differences between protein sequences, even when their overall sequence similarity is high.

8. ***DeepECTransformer***

DeepECTransformer [11], developed by the authors of the DeepEC model in 2023, represents a significant advancement in EC number prediction by employing the Transformer layers [12] as the neural network architecture. This model harnesses the power of self-attention mechanisms to capture complex dependencies within protein sequences, allowing it to process entire sequences holistically and identify critical features for accurate enzyme classification. By embedding protein sequences into a continuous vector space and utilizing multiple layers of self-attention and feedforward networks, DeepECTransformer focuses on the most relevant sequence regions, filtering out extraneous information. Like DeepEC, BLASTp is also integrated into the DeepECTransformer.

9. ***EnzBert***

Another Transformer-based model, introduced in 2023, is EnzBert [13]. EnzBert is based on a pre-trained language model named ProtBert-BFD [14]. ProtBert-BFD is the BERT architecture [15] pretrained on Big Fantastic Database (BFD) dataset [16] containing 2.1 billion protein sequences. EnzBert is the finetuned ProtBert-BFD on different datasets including SwissProt (release 2021_04). Model's parameters are shown in Table S2.

10. ***ProteinBERT***

ProteinBERT [17] is a BERT-based language model pretrained on UniRef90 protein sequences and their corresponding GO (Gene Ontology) annotations [18]. UniRef90 is a clustered database of protein sequences derived from the UniProtKB (Universal Protein Knowledgebase) and selected UniParc (UniProt Archive) sequences [3]. The model architecture is composed of two interconnected components: a local part that processes protein sequences and a global part that handles the GO annotation vector associated with those sequences. These components are pretrained simultaneously, enabling the model to effectively manage sequences of varying lengths, including those of extended size. Additionally, the model's computational and memory requirements increase linearly with sequence length, making it scalable. ProteinBERT can be fine-tuned for a variety of classification tasks, ranging from protein structure prediction to the prediction of biophysical properties. Model's parameters are shown in Table S3.

### Implementation

The selected models have their code available. However, DeepEC, ECPred, and DeepECTransformer do not provide their model training code, so we used their trained models instead of training them from scratch. Consequently, there is a possibility that some protein sequences from the test data were seen during the training of these models. The remaining models were trained from scratch for both the 30% and 100% similarity thresholds. The only model requiring pretraining was ProteinBERT. The pretraining and fine-tuning/training were conducted using each model's provided code. EnzBert also has a pretraining step, however, we skipped pretraining because it was expensive in terms of computation and time. ProteinBERT was pretrained and fine-tuned on a single NVIDIA Tesla A100-40GB GPU within a local cluster, named Alderaan[1], over the course of three weeks. The training and testing of other models were conducted on a single NVIDIA RTX 6000-40GB GPU. We alternated between these two clusters due to restrictions on library installations and storage limitation. Except for the homology-based models, all other models were implemented in Python. We have gathered their codes and library requirements into a unified benchmarking platform, available at https://github.com/dsaeedeh/EC-Bench.

### Setting of ensemble models

We considered two ensemble models in EC-Bench: majority voting and stacking.

1. ***Majority voting setting***

   For each input sequence, we collect the predictions from all models that generate at least one EC number, and the EC number with the highest frequency is selected as the final output. In cases where each predicted EC number has a frequency of 1, a random EC number is chosen as the final output. This method is particularly robust when the models tend to agree on the correct prediction, as it effectively reduces the impact of individual model errors by reflecting the consensus among the models.

2. ***Stacking setting***

   In our benchmarking task, we use stacking to leverage the strengths of different models by training a meta-model on the predictions of the base models. The base models first generate their predictions independently, and these predictions are then used as input features for the meta-model, which makes the final prediction. This approach allows us to capitalize on the complementary strengths of different models, potentially enhancing the overall predictive performance. The meta-model is carefully selected and trained on a validation set to ensure that it effectively combines the outputs of the base models.

   To stack models, we selected a few top-performing models to enhance the overall performance of the stacked model while also reducing training time. The selected models include DeepECTransformer, ECRECer, BLASTp, CLEAN, EnzBert, and ProteinBERT. The meta-model used for stacking is an instance of the MLPClassifier from the scikit-learn (sklearn) library [17]. The MLPClassifier, which stands for Multi-Layer Perceptron Classifier, is a type of neural network commonly employed for classification tasks. Parameters used to train the MLPClassifier is listed in Table S1.

   For training the meta-model, we utilized validation data from the Swiss-Prot 2023-02 release, ensuring consistency by applying the same preprocessing steps that were used for the training data. The validation dataset consists of 19,007 sequences for the 100% similarity threshold and 18,279 sequences for the 30% similarity threshold. After evaluating the models on this validation set and train set, we merged the output from training and validation data to train the meta-model, providing it with a comprehensive and diverse dataset to learn from. This approach aimed to maximize the meta-model's ability to generalize across different EC number predictions by incorporating a wide range of sequence variations during training.

[1] https://ccm-docs.readthedocs.io/en/latest/alderaan/
